# Playing the piano with a robotic third thumb: Assessing constraints of human augmentation

**DOI:** 10.1101/2020.05.21.108407

**Authors:** Ali Shafti, Shlomi Haar, Renato Mio, Pierre Guilleminot, A. Aldo Faisal

**Affiliations:** Brain and Behaviour Lab: Dept. of Bioengineering, Imperial College London, SW7 2AZ, London, UK; Dept. of Computing, Imperial College London, SW7 2AZ, London, UK; Behaviour Analytics Lab, Data Science Institute, SW7 2AZ, London, UK; Dept. of Brain Sciences and UK Dementia Research Institute – Care Research & Technology Centre, Imperial College London, W12 0BZ, London, UK; UKRI CDT in AI for Healthcare, Imperial College London, SW7 2AZ, London, UK; MRC London Institute of Medical Sciences, W12 0NN, London, UK

## Abstract

Contemporary robotics gives us mechatronic capabilities for augmenting human bodies with extra limbs. However, how our motor control capabilities pose limits on such augmentation is an open question. We developed a Supernumerary Robotic 3rd Thumbs (SR3T) with two degrees-of-freedom controlled by the user’s body to endow them with an extra contralateral thumb on the hand. We demonstrate that a pianist can learn to play the piano with 11 fingers within an hour. We then evaluate 6 naïve and 6 experienced piano players in their prior motor coordination and their capability in piano playing with the robotic augmentation. We show that individuals’ augmented performance could be predicted by our new custom motor coordination assessment, the Human Augmentation Motor Coordination Assessment (HAMCA) performed pre-augmentation. Our work demonstrates how supernumerary robotics can augment humans in skilled tasks and that individual differences in their augmentation capability are predictable by their individual motor coordination abilities.

## Introduction

From ancient myths, such as the many-armed goddess Shiva to modern comic book characters, augmentation with supernumerary (i.e. extra) limbs has captured our common imagination. In real life, Human Augmentation is emerging as the result of the confluence of robotics and neurotechnology. We are mechatronically able to augment the human body; from the first myoelectric prosthetic hand developed in the 1940s ^1^ to the mechanical design, control and feedback interfaces of modern bionic prosthetic hands ^e.g. 2–5^. Robots have been used to augment the bodies of disabled humans, restoring some of their original capabilities ^e.g. 6–9^. Similar setups can augment healthy users beyond their capabilities, e.g. augmenting workers in industrial settings through intelligent collaboration ^e.g. 10–12^, or equipping them with additional arms to perform several tasks concurrently ^e.g. 13,14^. The latter fits within a particular area of human augmentation robotics which is referred to as supernumerary robotics. These are robotic systems, typically worn by the user, to extend their body and its physical capabilities. However, a major question is, to what extent does human motor control have the capability of adapting and learning to use such technologies efficiently ^15^, and can it be predicted from human motor coordination. The supernumerary and augmentative nature of this area of research presents an interesting challenge on how to map human motor commands to robot control.

Supernumerary robotic limbs ^e.g. 16^ are envisioned to assist human factory workers, and adapted for different types of applications ^e.g. 14^. The introduction of supernumerary robotic fingers developed as a grasp support device ^17^ led to further exploration on optimal materials and mechanical designs for supernumerary robotics ^e.g. 9,18,19^. Supernumerary robotic fingers have been particularly envisioned, and successful in grasp restoration for stroke patients ^e.g. 9,20,21^. However, regardless of the mechanism, material, and use case, given the presence of a human within these devices’ control loops, the control interface is of essential importance.

It was recently shown that for polydactyly subjects who possess six fingers on their hands, the control interface involves a cortical representation of the supernumerary finger ^22^. Unlike polydactyly, supernumerary robotic limbs and fingers must utilize indirect control interfaces to achieve the same goal – to enable more complex movements and better task performance. Abdi et. al investigate the feasibility of controlling a supernumerary robotic hand with the foot ^23^. Others have focused on electromyography (EMG) as the control interface – both in supernumerary robotic fingers ^e.g. 9^, and supernumerary robotic limbs ^e.g. 24^. Other interfaces used for supernumerary robotics include inertial measurement units ^e.g. 21^, voice ^e.g. 25^, pushbuttons ^e.g. 18^, and graphical user interfaces ^e.g. 26^. Researchers have also explored indirect control interfaces, e.g. using the concept of grasp synergies ^27^ to assume that the supernumerary robotic finger’s posture will be highly correlated with that of natural fingers during manipulation, allowing supernumerary robotic finger control through natural movement of existing fingers ^28^. Importantly, all these user interfaces focus on the interface and not the user.

While extensive research has been conducted on the mechanical design, interface, and control of supernumerary robotics, there is a gap in understanding the role of human motor control in the success and adoption of these robotic human augmentation systems. In the rapid development of human augmentation little attention is devoted to how humans interact with the technology and learn to control it ^15^. There are clear needs for neuroscience and robotics research to come together in analysing such scenarios ^e.g. 29^. Non-invasive control interfaces for supernumerary robotics typically rely on substitution, i.e. using the movement of a natural human limb for the control of the augmentation robot ^e.g. 30^. Learning to control a supernumerary robot limb or finger thus becomes a complex process which involves learning to utilize one movement (set muscles activations) to perform a new movement. The field of motor neuroscience has extensively studied the control mechanisms and learning processes of perturbed movements. Some research paradigms directly address this augmentation challenge as they utilize arm movement in one direction to move a cursor on the screen in a different direction, accounting for a rotation perturbation ^31–34^ or a mirror reversal perturbation ^e.g. 35–37^. In these settings, one can predict subjects’ learning from the task-relevant variability in their unperturbed movements ^e.g. 38,39^. Nevertheless, these studies were done on simplistic lab-based tasks and only recently the field is starting to address the complexities of real-world movement and to ask to what extent those lab-based findings generalize to real world motor control and learning ^40,41^. While the relationship between task-relevant variability and learning performance seems to generalize to real-world tasks, defining task relevance is less trivial ^e.g. 40^ and the learning mechanism can differ between users ^e.g. 42^.

In human performance research, such as sports science and rehabilitation, there are significant efforts to predict future performance. In sports science, there is an attempt to predict athletes’ future success for talent identification purposes. Motor coordination and motor learning are often used as predictors ^e.g. 43–46^. Similar approaches are used in rehabilitation research to predict skill learning capacity following traumatic brain injury, stroke, or neurodegenerative disease ^e.g. 47,48^. Here we are looking into the predictability of future performance with augmentation technology. We specifically ask which aspect of motor coordination is a better predictor of performance in the same task with and without the device.

To address this, we have created an experimental setup to study how different parameters within the remit of human motor control contribute to successful control, coordination and usage of a human augmentative robotic system, using a set of motor coordination tests that we developed: the Human Augmentation Motor Coordination Assessment (HAMCA). We have created a 2 degrees of freedom (DOF) robotic finger, worn on the side of the hand, to augment human finger count to 11, effectively giving them a 3rd thumb. We call this the supernumerary robotic 3rd thumb (SR3T – engineering and design previously described in ^49^), and we study its usage in a skilled human task: playing the piano. The piano is a setting which involves the use of all fingers of the hand, and hence a good environment to consider for testing the augmentation of fingers. Furthermore, piano playing is structured both in spatial and temporal dimensions, allowing for quantification of the performance in both aspects.

## Results

We developed a mechanically powerful supernumerary robotic 3rd thumb (SR3T) and means for interfacing it with human users, initially through a combination of foot and thumb motions, directly controlling the two degrees of freedom of the SR3T. We then tested an experienced piano player in an unconstrained pilot experiment, allowing them to freely play the piano and improvise while wearing the SR3T. We observed that within 1 hour of playing the piano, the participant incorporated the SR3T in their piano playing, effectively playing the piano with 11 fingers (see Fig. 1G-H, and supplementary video). Based on this outcome, and feedback from the participant, we upgraded the control interface to be solely based on foot motions (see Fig. 1, and Methods), for more robust control, and to limit the interface to one limb. We then set out to understand the constraints affecting success with the SR3T, by devising protocols and behavioural markers for motor coordination evaluation: the Human Augmentation Motor Coordination Assessment (HAMCA). We also developed a piano sequence playing task as well as measures for assessing the quality of playing. Finally, we systematically evaluated the SR3T on human subjects, and predicted how well subjects would be able to perform in playing the piano sequence with an augmented additional finger, based on the basic motor coordination assay from the HAMCA. Twelve right-handed participants (6 experienced pianists and 6 naïve players), attended 2 experimental sessions held on separate days in the lab. In the first session they performed the HAMCA set of 8 tasks to assess their hand and foot motor coordination. This set was developed to investigate the possibility of a priori prediction of how well each individual human user can learn to use an augmented device. From the HAMCA we extracted 8 scores as measures of motor coordination within hand and foot motions (see Methods). In the second session the subjects learned to play a sequence on the piano and then repeated it with our human augmentation device, the supernumerary robotic 3rd thumb (SR3T), operated through foot motions as the interface (Fig. 1).

**Fig. 1.**
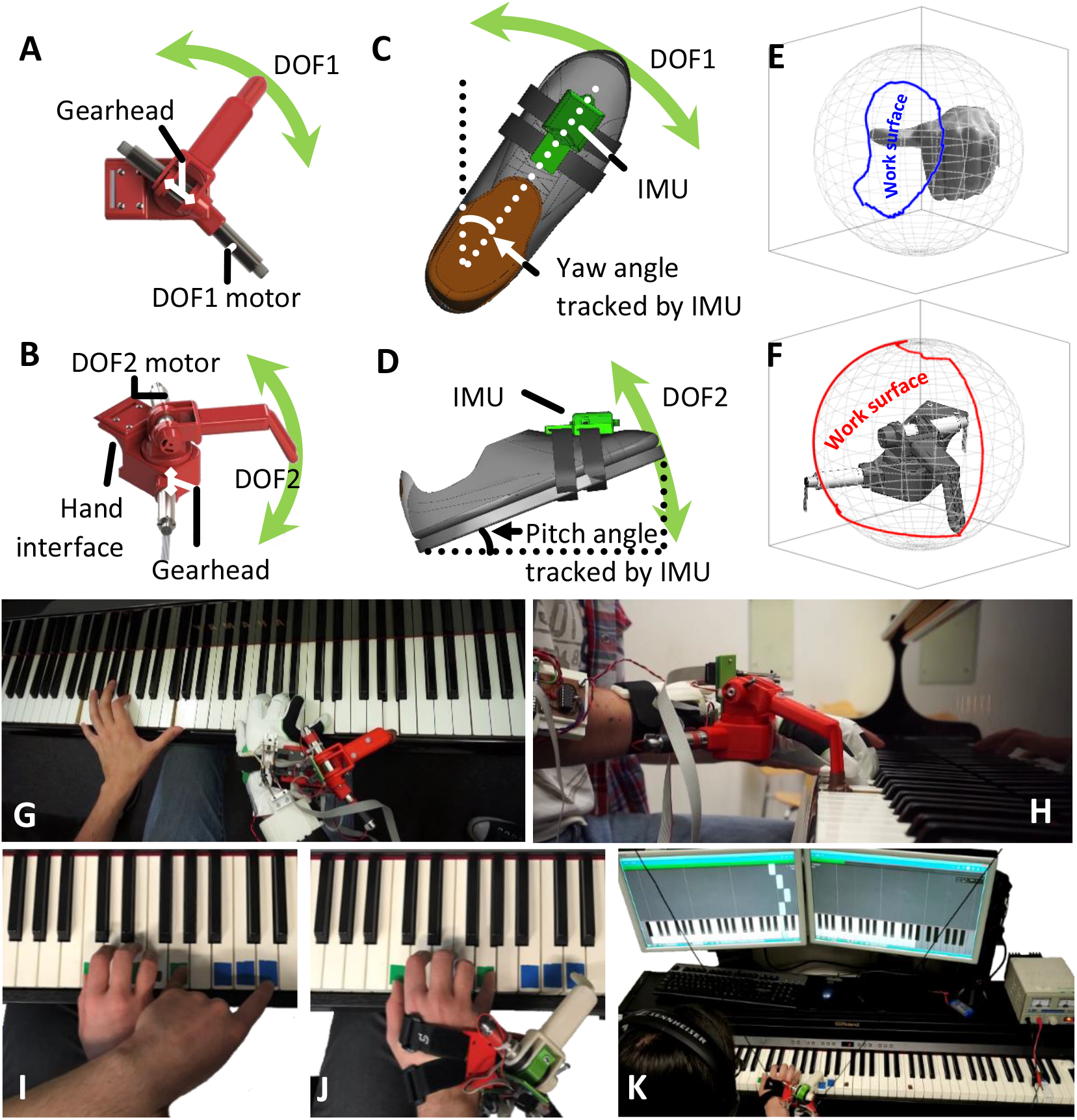
Piano playing task setup. **(A)** Top view rendering of the SR3T, showing the horizontal motion DOF and relevant motor. **(B)** Side view rendering of the SR3T showing the vertical motion DOF and relevant motor. **(C)** Top view rendering of the SR3T control interface for the 1st degree of freedom (DOF); the participant controls the motion of the SR3T using their right foot, captured through an inertial measurement unit (IMU) worn on the foot. **(D)** Side view rendering of the SR3T control interface for the 2nd DOF. **(E)** Work surface of a human thumb end-point projected on a sphere for comparison with **(F)** the work surface of the SR3T end-point projected on a sphere – augmenting work surface range for the human (see methods). **(G, H)** Top and side view of the unconstrained pilot experiment: an experienced piano player freely improvising on the piano while wearing and making use of the SR3T, effectively playing 11-fingered piano within 1 hour of use. **(I)** Systematic experiments: playing the piano sequence using 5 fingers of the right hand plus the left-hand index finger (LHIF) and **(J)** Playing the sequence using the SR3T. **(K)** A participant plays the sequence of notes as displayed on the monitors in front of them, using the SR3T.

To give a sense of the HAMCA data, distributions of scores, and to highlight any significant differences between groups we present a summary of results in Fig. 3. The hand and foot motor-coordination scores from the HAMCA, recorded during the first experimental session showed moderate differences between the pianists and the naïve players (Fig. 3A). We took a conservative approach to capture even marginally significant group differences and thus did not adjust for multiple comparisons. Yet, in a permutation test there were no significant group differences in any of the piano-based tasks (Piano Jumping, Piano Timing, and Piano Loudness). The only measure where the pianists performed marginally significantly different to the naïve players was Hand Dimensionality (p = 0.017), which is based on the assembly of toys (see Methods). On the other hand, in the Foot Up-Down Spatial measure the naïve players showed higher scores than the pianists (p = 0.013), though we believe this difference is possibly the result of a sample bias due to the small N. In both groups the inter-subject variabilities were relatively evenly distributed.

The correlation matrix between the subjects’ motor-coordination scores (Fig. 3B) suggests relatively weak dependencies, i.e. a subject who showed high coordination in one task did not necessarily show high coordination in any other task. Even without multiple comparisons correction there were no significant dependencies within the foot measures and the only dependency within the hand measures was between the Piano Jumping and Piano Timing scores (r = 0.64, p = 0.026). The foot and hand timing scores were correlated (Foot Up-Down Temporal and Piano Timing: r = 0.68, p = 0.015; Foot Up-Down Temporal and Piano Jumping: r = 0.66, p = 0.020). These three tasks are metronome based, thus measuring rhythmic coordination. Lastly, we found a correlation between two movement precision tasks: Foot Balance and Piano Loudness scores (r = 0.61, p = 0.035).

In the second experimental session, all subjects performed 10 trials of the Piano Playing task, using their left-hand index finger (LHIF) to play notes further to the right of their right-hand. This was followed by an additional 10 trials of Piano Playing with the SR3T, where subjects use both degrees of freedom within the SR3T for the horizontal reach for the notes to the right and the vertical motion to play the note. Each note is scored as 1 if played correctly and perfectly on time, with the score linearly decreased to 0 if played half a beat late or early. Incorrect notes are also scored as 0. Note scores are averaged to report a final piano playing score (see Fig. 4B, and Methods). In both tasks (playing with the LHIF and with the SR3T) subjects showed improvement over the first 5 trials after which they plateaued (Fig. 4, right). Thus, for all future analysis we averaged over trials 5 to 10 to have a single piano playing score for each of these tasks. Here as well, there were no significant differences between the pianists and the naïve players in any of the trials played with their LHIF (t-test p > 0.12) and with the SR3T (t-test p > 0.32). As performance varied randomly in each subject over the plateau, we also pulled all plateau trials together and tested again for group differences. Testing over all plateau trials (5-10), pianists were significantly better in playing with their LHIF (t-test p = 0.017) but there were no group differences in playing with the SR3T (t-test p = 0.9). Therefore, we merged the two groups, and all further analysis was done on all 12 subjects together.

The Piano Playing with SR3T score is our metric for performance with the human augmentation device. Augmentation here refers not necessarily to improvement in performance, but rather an augmented reach, allowing to play notes that are further away, without moving the hand or using another hand (the case in LHIF). The fundamental question is to what extent can the SR3T score be explained by motor-coordination measures. The correlations between the subjects’ scores in the Piano Playing tasks and the motor-coordination measures suggest different dependencies for playing with and without the SR3T (See supplementary, Fig. S1 & Fig. S2). The scores in the Piano Playing task with the LHIF were correlated with Foot Tracking and Foot Up-Down Temporal scores (r = 0.66, p = 0.019 and r = 0.64, p = 0.026, respectively). The scores in Piano Playing with SR3T were correlated only with the Piano Loudness scores (r = 0.59, p = 0.044). We further investigated the correlations between Piano Playing with SR3T and the motor-coordination scores with Spearman rank correlation scores (Fig. S1). Foot Up-Down Temporal was the only measure which showed marginally significant Spearman correlation with the Piano Playing with SR3T score (r(Spearman) = 0.67, p = 0.02). The Piano Playing scores without and with SR3T were correlated (r = 0.63, p = 0.028), and even better correlated with Spearman rank correlation (r(Spearman) = 0.71, p = 0.012).

We then asked which combination of HAMCA motor-coordination scores can better explain the LHIF and SR3T scores. We considered all possible combinations of the HAMCA features, and different number of features being selected. We fit linear models to all given combinations, trying to explain the piano playing scores with the LHIF and SR3T. To compare between these models of different complexity we used the corrected Akaike information criterion (AICc) which is modified for small sample sizes. AICc estimates the amount of information that is lost while fitting a model and thus can measure the quality of different models relative to each other. Results for AICc and the correlation between the real score and the predicted score (r) can be seen in Fig. 5A for the LHIF and Fig. 5B for the SR3T, for all combinations up to 5 features (higher number of features result in increasingly deteriorated models), with the best models and their selected features highlighted. When the sample size is small, information criteria might be biased to select models with more parameters and overfit, the AICc was developed to address such potential overfitting. With this correction, the AICc suggests that the best model for predicting the LHIF score is a 2-feature one, whereas for the SR3T, a 4-feature model would perform better. Looking at the selected features, in the case of the LHIF, both models show similar high performance, using Foot Tracking as one feature, with the second feature being Foot Up-Down Temporal in one model (AICc = −28.37, r = 0.86, p = 0.0023) and Piano Jumping in the other (AICc = −27.81, r = 0.85, p = 0.0028). The best performing SR3T model uses Foot Up-Down Spatial, Foot Up-Down Temporal, Piano Timing and Piano Loudness (AICc = −16.66, r = 0.95, p = 0.0012).

## Discussion

In this study we examined the role of human motor coordination in the successful adoption of human augmentation technology. We had previously described the different neurocognitive barriers to successful embodiment and use of robotic augmentation devices ^15^. Here, following the operational definition set out in the same work ^15^ for the embodiment of robotic augmentation as the ability to use extra limbs in natural tasks, we focused on how different parameters within the remit of human motor control contribute to successful control of a supernumerary robotic finger in an augmented piano playing task. We created a supernumerary robotic 3rd thumb (SR3T), controlled through substitution, initially with a combination of the natural thumb and the foot wearing different sensing modalities. We first demonstrated, in an unconstrained pilot experiment, the feasibility of human augmentation with the SR3T, with one experienced piano player using the device to freely play the piano using 11 fingers, within 1 hour of wearing it. We then updated our SR3T interface, so that both degrees of freedom are controlled through a single IMU worn on the foot, and developed the HAMCA set to assess the participants’ motor coordination prior to piano playing. This is followed by our subjects playing a piano sequence on the piano using their natural fingers, and then using the SR3T along with their natural fingers. Our HAMCA metrics can be used to predict and explain both the task performance with, and without robotic augmentation. While the LHIF task can be explained by two features relating to timing precision, the SR3T task requires four features, involving both timing and movement precision. This suggests that performing the task with robotic augmentation, requires additional levels of complexity.

Our work shows the possibility of humans being able to quickly acquire a skilled behaviour, such as playing piano sequences, with a human augmentative robotic system. Both naïve piano players (i.e. without prior experience) and piano playing experts demonstrated the same ability to integrate the supernumerary robotic limb into playing simple piano sequences: We saw no group differences in the performance with the SR3T, suggesting that integrating robotic augmentation is primarily driven by a priori motor coordination skills and not affected significantly by expert motor domain knowledge. We designed our piano piece to ensure comparability with the experimental setup, so that pre-augmentation the participants would play the main notes with their right hand and then play the additional notes with the left hand index finger, therefore limiting us to one-handed pieces. The structure of the music piece itself also needed to be simple enough so that piano-naïve participant could acquire it within a reasonable amount of time and not immediately fail. Further experiments are needed to examine whether pianists’ expertise will give them an advantage in the augmented task, for more complex piano sequences.

There were also not many significant differences between the two groups within the HAMCA scores. This might be due to the design of the tasks within HAMCA not capturing that difference. Importantly, unlike studies of piano skills that investigate different motor performance metrics in piano players ^e.g. 50,51^, HAMCA was developed to capture basic motor control and coordination to predict augmentation. We used the task space (piano key presses) and the interface space (foot control) but did not design motor performance metrics in piano playing. Hence, perhaps it is not surprising that the task-space metrics were not performing as such, evident by the lack of difference between pianists and naive players.

In the piano playing task, all subjects (naïve players and experienced pianists alike) initially showed improvement in accuracy from trial to trial (i.e. learning). This was a short learning process which plateaued quickly after 5 trials. For all subjects the plateau with the SR3T was significantly lower than with their LHIF, which is to be expected, particularly in early stage of SR3T use. This fast learning within a session and low plateau (which leaves much room for improvement in future sessions) are hallmarks of early motor skill learning. This is in line with many evidence of multiple time scales in skill learning where fast improvement in performance occurs in the initial training and plateau within a session, and slow improvement develops across sessions ^e.g. 52–55^. Accordingly, learning to play the piano, augmented with the SR3T, seems to be a novel motor skill learning task. Further support can be found in the group differences while playing the piano with and without the SR3T. When subjects played with their own LHIF, across all plateau trials pianists performed significantly better than naïve players, as expected based on their piano experience, though there were no significant group differences on a trial by trial basis, not during learning nor during plateau. When subjects played with the SR3T there were also no significant group differences on a trial by trial basis, but also across all plateau trials pianists did not perform better than naïve players. This suggests that playing augmented with the SR3T is not a trivial extension of the regular piano sequence playing task with your own finger, but a novel skill that the subjects need to learn.

To further interrogate the novelty and complexity of the augmented task, we explored the differences in the way combinations of motor coordination measures from HAMCA can predict performance with the SR3T and the LHIF. Our results show that the SR3T score is best explained with twice as many features as required for the best model for the LHIF, suggesting a higher complexity in the augmented task. Furthermore, the HAMCA features selected for the top LHIF models mainly measure timing precision (Foot Tracking paired with Foot Up-Down Temporal or Piano Jumping), whereas the best SR3T model uses two timing precision features (Foot Up-Down Temporal and Piano Timing) together with two movement precision ones (Foot Up-Down Spatial and Piano Loudness), see Fig. 5. This suggests that the SR3T model, relating to the augmentation task, needs to capture not just task-related features through timing precision, but also movement precision features relating to both the interface (Foot Up-Down Spatial) and the task (Piano Loudness).

It is important to consider the meaning of these results in the context of prosthetics. Prosthetics replace a limb that was lost whereas with the SR3T, and with supernumerary robotics in general, the human is operating a new, additional limb – in our case a thumb. Our augmentation is done through substitution, i.e. we use an existing limb to operate an additional one. While many active upper limb prosthetics with non-invasive interfaces rely on EMG from residual muscles^56–58^, there is still interest in movement-based control due to challenges with EMG interfaces^59,60^. Foot motions in particular, detected through various sensing modalities, have been used as prosthetic interfaces^61–64^. While researchers investigate various important factors in prosthesis choice and success, such as cognitive, social, or physical ones ^e.g. 65–67^, in an effort to tackle the ongoing adoption problem facing modern prosthetics^68,69^, there is less focus on the role of the end-user’s motor coordination, which we argue is particularly important in substitutional interfaces. Our results show that motor coordination metrics are highly predictive of success in substitutional augmentation. We have argued before that not accounting for such factors, can lead to barriers in adoption of technology^15^.

Previous work on interfaces for supernumerary robotics have shown that the foot can generally serve as a good interface for robotic limbs working collaboratively with the user’s hands ^e.g. 23,70^. Abdi et al. ^23^ study the control of a third robotic hand via the foot in virtual reality, for robotic surgery applications, showing similar learning trends to what we observe here. Similar learning within a session was also reported with another foot interfaced third thumb^30^. We see similar effects where roboticists have used legs and feet as a multi-DOF control interface for successfully teleoperating two ^71^ or four ^72^ robotic arms, albeit in less skilled tasks than what we show here. Saraiji et. al. show that subjects significantly increased their self-reported sense of embodiment of the tele-operated robots over the course of the experiments, i.e. 40 minutes ^71^. Results obtained with adaptive foot interfaces for robot control ^e.g. 73^, where data-driven approaches are used to create subject-specific motion mapping, are in line with our findings. Huang et al. ^73^ report that inter-subject variability decreases once a subject-specific motion mapping is enabled. This confirms the dependency of robot control performance on metrics inherent to each subject, which we present here to be a combination of motor coordination metrics relating to the task and the robotic interface.

We show here that substitutional interfaces make the system reliant on human motor coordination skills. We also demonstrate the possibility of applying substitution across different levels of the biomechanical hierarchy. The foot, which in terms of the biomechanical hierarchy is equivalent of the entire hand, is used here as the interface-space controlling a thumb, which is further down the biomechanical hierarchy. These results sit at one end of the spectrum of solutions for controlling an augmentative device, which goes from substitution all the way to direct augmentation via higher level control, either brain-machine-interfacing or cognitive interfaces such as eye gaze decoding. We previously showed that the end-point of visual attention (where one looks) can control the spatial end-point of a robotic actuator with centimetre-level precision ^8,74,75^. This direct control modality is more effective from a user perspective than voice or neuromuscular signals as a natural control interface ^76^. We showed that such direct augmentation can be used to control a supernumerary robotic arm to draw or paint, freeing up the two natural arms to do other activities such as eating and drinking at the same time ^13^. But such direct augmentation has to date not achieved augmenting fine motor skills such as playing the piano, as playing this instrument requires not just the execution of a note: it is not a simple button-press exercise but requires fine grade expression of temporal and spatial motor coordination across robotic and natural fingers. We show that we can predict the degree to which subjects can integrate supernumerary limbs into their natural body movements, as a function of their basic motor skills. Thus, our work shows that we can achieve effective augmentation but also predict the capability of individuals to embody supernumerary robotic limbs in real-world tasks, which has impact for robotic augmentation from healthcare to agriculture and industrial assembly e.g. in the aerospace industry.

## Materials and Methods

### Experimental design

For our unconstrained pilot experiment, a right-handed piano player was selected to wear the SR3T and freely improvise. For our systematic follow-up experiments we developed a set of measurement protocols and behavioural biomarkers, the Human Augmentation Motor Coordination Assessment (HAMCA), and ran this set of tests to assess hand and foot coordination (due to the foot being the control interface for our robotic system, described below under *Setup*) and piano-related skills. As opposed to existing motor assessments such as the Purdue pegboard ^77^, the motor domain of the NIH Toolbox ^78^, the Jebsen-Taylor hand function test ^79^ or the Action Research Arm Test (ARAT) ^80^ among others, which tend to be focused on dexterity, and are mainly used to quantify the extent or progress of motor disabilities, here we are interested in assessing specific human motor coordination aspects which relate to the interface-space (foot) and task-space (hand use over the piano) of our piano playing task. The HACMA set includes both spatial and temporal evaluations. From these coordination tasks we extracted 8 hand and foot motor-coordination scores. Finally, the participants were given specific melodies to play on the piano with and without the SR3T. The melody was designed to require 6 fingers, forcing the participant to either use their left-hand index finger (LHIF), or the SR3T if they are wearing it. Table 1 summarises the experimental setup procedure and how they map to results.

**Table 1.** Experimental procedure

| First Session – 2 hours | Second session – 1 hour |  |
| --- | --- | --- |
| Foot Balance (15 trials) – 20 mins | Piano Playing with LHIF – 25 mins |  |
| Foot Up-Down (15 trials) – 15 mins | 5 Practice Trials | 10 trials recorded at 80bpm (last 5 count for score) |
| Foot Tracking (6 trials) – 10 mins |  |  |
| Piano Timing (25 trials) – 15 mins | SR3T setup and calibration – 10 mins |  |
| Piano Jumping (15 trials) – 10 mins | Piano Playing with SR3T – 25 mins |  |
| Piano Loudness (25 trials) – 15 mins | 5 Practice Trials | 10 trials recorded at 80bpm (last 5 count for score) |
| Hand Dimensionality (2x toy tasks) – 35 mins |  |  |

### Subjects

Twelve right-handed participants took part in our systematic experiments (mean age 23.3+/−2.8 years). Six of the participants had played the piano for several years (pianists group) and the other six did not have any piano playing experience (naïve group). All of the participants from the pianists group had at least 5 years of piano training (range 5-21 years, mean 10.6+/−5.4 years). Two participants of the naïve group had over 5 years’ experience of guitar playing. All participants gave informed consent prior to participating in the study and all experimental procedures were approved by Imperial College Research Ethics Committee and performed in accordance with the declaration of Helsinki.

### Setup

We created an experimental setup to investigate how individual motor skills contribute to the performance of a human user of a supernumerary robotic thumb; i.e. a robotic augmented human. To this end, we have created a 2 degrees of freedom (DoF) robotic finger that users can wear on the side of their hand, effectively augmenting them with a third thumb. The design, creation and initial testing of the supernumerary robotic third thumb (SR3T) was reported in ^49^, and is the same setup used for our unconstrained pilot experiment with a single piano player participant. The design specifications for the SR3T were derived from the design requirements of a fully spherical operating thumb ^81^ and the natural eigenmotions of human thumbs in daily life activities ^82^. The SR3T is attached to the user’s right hand and is controlled through the user’s right foot. In our original implementation, used for the pilot experiment, the vertical motion of the foot was measured using an accelerometer, together with horizontal motion data obtained with a flex sensor worn on the natural thumb on the augmented hand, to control the vertical and horizontal DoFs of the SR3T, respectively ^49^. For our main experiments, we updated the interface, using a 9DoF inertial measurement unit (IMU - Bosch BNO055, breakout board by Adafruit) for increased stability, and to limit the interface to a single limb, i.e. the foot. The unit can provide absolute orientation measurements (with respect to the earth’s magnetic field) thanks to an onboard sensor fusion algorithm. Absolute orientation can then be extracted as Euler vectors, at 100Hz. In this setup, the SR3T’s two DoFs correspond to horizontal and vertical movements of the robotic fingertip. These are mapped to horizontal and vertical movements of the foot, i.e. yaw and pitch, respectively. Once the subject is wearing the SR3T on their hand, and the IMU on their foot, they are asked to sit with their foot on the ground facing the piano. The SR3T is moved horizontally for the fingertip to face the forward position as well. The values read by the IMU for the orientation of the foot, and by the motor encoders for the position of the SR3T are recorded. The subject is then asked to rotate their foot clockwise, with the heel as the centre of rotation, to their maximum comfortable reach (typically 45 degrees from the forward-facing pose). The SR3T fingertip is also moved accordingly, to the maximum horizontal position on the right, and values recorded as before. These are used to map the horizontal motion of the foot to that of the SR3T, with a similar process for vertical motions. The setup can be seen in Fig. 1.

In order to investigate the workspace augmentation achieved by the SR3T optical markers were placed at the tip and base of the SR3T, with the SR3T then activated by the subject to move in its full range of motion. Similarly, optical markers were placed at the tip and base of the subject’s left-hand thumb with them performing the maximum range of thumb movement while the motion was optically tracked. We used three OptiTrack Prime 13W cameras with the Motive software for motion capture (NaturalPoint, Inc. DBA OptiTrack, Oregon, USA). The results can be seen in Fig. 1.E and F; the thumbs’ end-point surface is mapped onto a sphere, assuming the base of the thumbs are situated at the centre. Based on these measurements, the SR3T has an end-point work surface that is 4 times that of the human thumb. Furthermore, we used the same camera system to measure delay between motor intention and action, by placing markers on the user’s foot as well as the SR3T; the mean delay is measured as 85msec. This delay does not affect our piano playing scoring scheme (see below for details).

For the piano playing tasks and piano related hand coordination tasks we used a digital piano (Roland RP501R-CB, Roland Corp., Osaka, Japan). The piano was connected to a PC with a MATLAB script establishing communication through its MIDI interface. Each keystroke on the piano was received by the MATLAB script as a MIDI message which comprised data regarding the note played, time of keystroke (with a 1ms resolution) and the keystroke velocity, which leads to proportional loudness of the note played.

#### HAMCA foot coordination tasks

##### Foot balance task

A Wii Balance Board (Nintendo Co. Ltd., Kyoto, Japan) together with the BrainBloX software ^83^ was used. The board (Figure 2A) is made of four pressure plates and the software interface displays the real-time centre of pressure computed by the Wii Balance Board across all four plates, and relative to the board’s coordinate system.

**Fig. 2.**
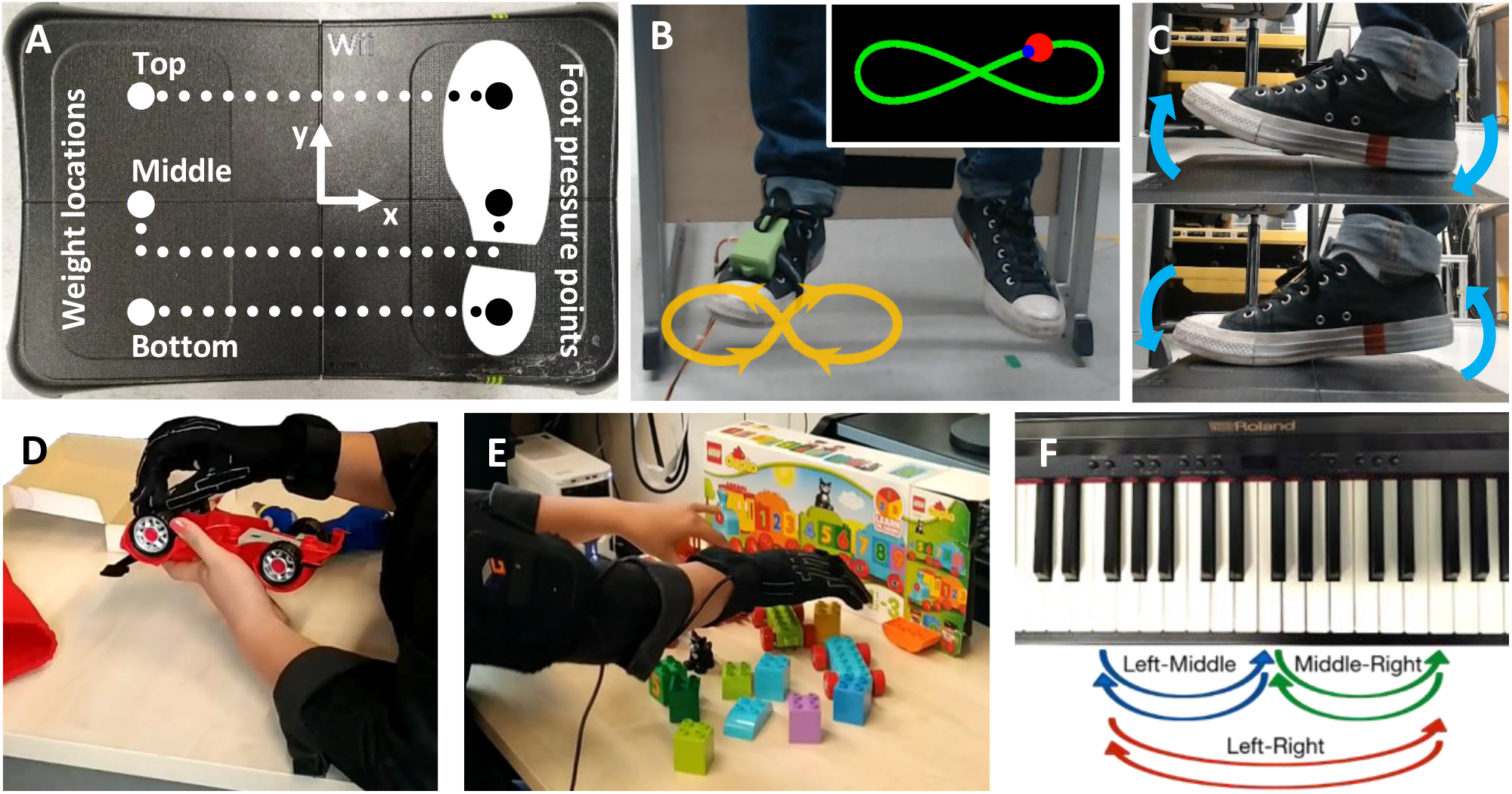
Human Augmentation Motor Coordination Assessment (HAMCA) – a set of simple behavioural tasks to predict the ability of human augmentation (see Methods for details). **(A)** Balance board force measurement platform for the Foot-balance task. **(B)** Foot motion trajectory (curve and arrows) during the Foot-tracking task required to move the foot in a figure of 8 in a plane perpendicular to the resting foot’s major axis. (Inset) visual feedback to the participant on the computer screen in front of them, showing the desired trajectory (green curve), the red dot indicating the desired location of the foot tip for pacing the foot movement and the current location of the foot tip (blue dot). **(C)** See-saw like foot motion in the sagittal plane of the foot during the Foot up-down task while seated. **(D,E)** Measurement of motor coordination complexity in the fingers of the right hand by assembly of a toy car and of a toy train in the Toy Assembly Task. **(F)** Piano jumping task: Piano key sequences to be played with individual fingers to capture hand and finger timing acuity when jumping between notes. Performance is assessed by timing and key board press down errors movement between 3 keys (spaced 1 octave apart) labelled “Left”, “Middle” and “Right”.

**Fig. 3.**
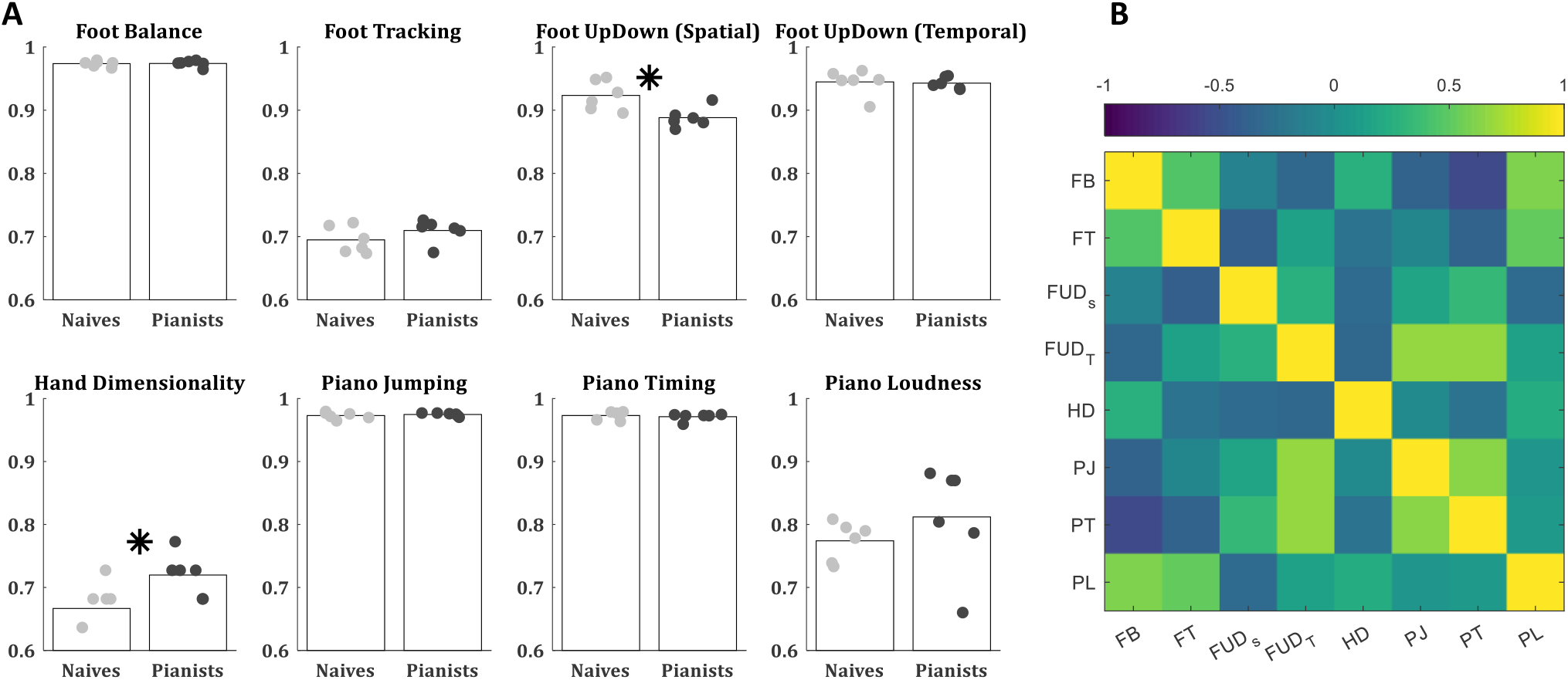
Performance of all subjects in *the HAMCA*. Showing results for 6 naïve (grey dots) and 6 experienced piano players (black dots). (A) Accuracy in HAMCA tasks (foot Balance - FB, foot tracking – FT, foot up down spatial – FUD_S_, foot up down temporal – FUD_T_, hand dimensionality – HD, piano jumping – PJ, piano timing – PT, piano loudness - PL). (B) Pearson’s correlations between the accuracies of all HAMCA tasks across subjects. The 6 naïve and 6 experienced players were lumped together as individual performances were not different between the groups (see main text).

**Fig. 4.**
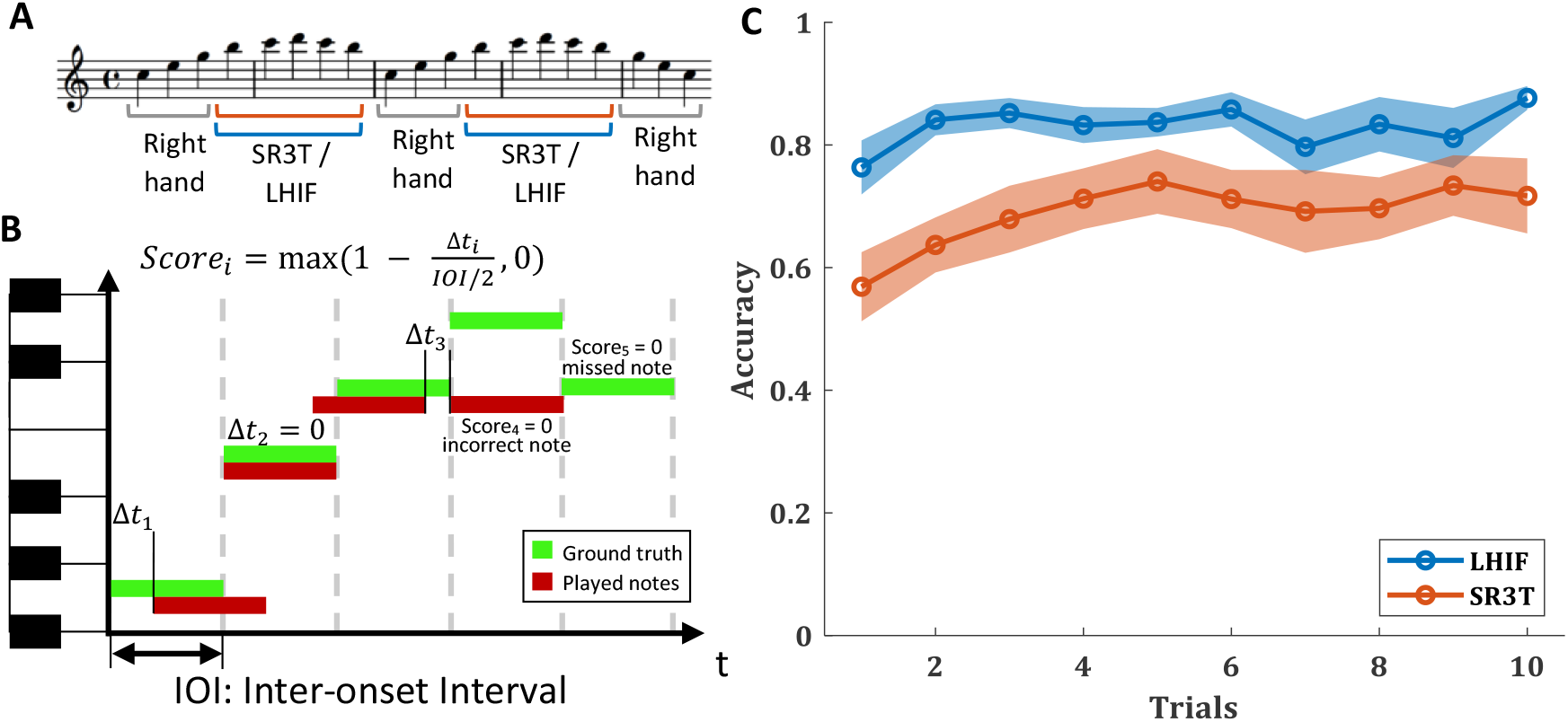
Piano playing performance. **(A)** The note sequence played for the piano playing task. Notes exclusively played with the right hand, and those with the SR3T or LHIF are marked, **(B)** Visualisation of how each individual note is scored linearly based on delay from the beat. Incorrect notes and skipped notes are assigned a score of 0, full sequence score is the average of all individual scores, **(C)** Accuracy over trials with the SR3T (orange) and without, using the LHIF (blue).

**Figure 5.**
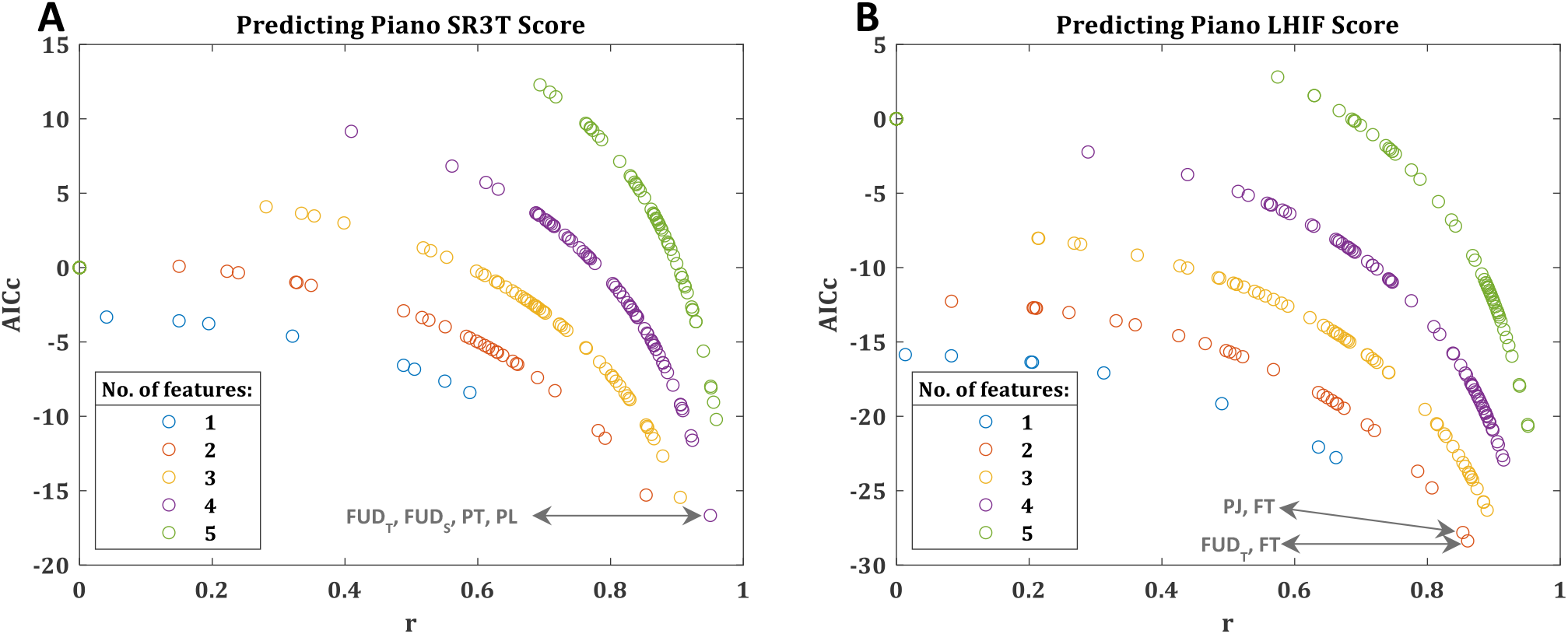
Model selection predicting LHIF and SR3T piano playing. All possible combination of different number of features being selected from the HAMCA metrics are considered and used to fit generalised linear models, with AICc and r reported. Model performances for all combinations up to 5 features shown, with best model and its features highlighted in **(A)** SR3T score prediction and **(B)** LHIF score prediction. Selecting more than 5 features results in increasingly deteriorating models.

Weight plates (70 N) were placed on the left side of the board, moving the centre of pressure away from the system origin. Subjects were seated on a chair, placing their right foot on the Wii Board, and then had to move the centre of pressure back towards the origin by applying pressure on the right side of the board with their right foot. The plates were placed in three positions (Figure 2A), with five trials per position, resulting in a total of 15 trials, performed in random order. Before the beginning of each trial, participants were asked to place the centre of pressure as close as possible to the origin. Once they stated their readiness and after a 3-seconds countdown, a 15-seconds recording was started. Samples were recorded at 85 Hz. The resulting motor-coordination score is computed according to equation (1):

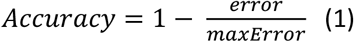

Where error corresponds to the mean Euclidean distance of the centre of pressure from the origin of the coordinate system across all recorded samples. The maximum error corresponds to the error computed if the subject was not acting on the platform.

##### Foot up-down task

The same setup as in the Foot Balance Task was used, without the weights. A steady beat was played with a metronome, which the subjects had to match while sitting on a chair with their right foot placed on the Wii Board, by moving their feet from a toe-lifted (dorsiflexion) to a heel-lifted (plantarflexion) pose and vice versa (see Figure 2C). The pressure exerted by the foot had to match an upper and lower target value marked on the screen. Ideally, the output should resemble a square signal with a period equivalent to that of the beat on the metronome. Subjects performed 15 trials in random order, five at each selected tempi: 40bpm, 60bpm and 80bpm.

Two types of motor-coordination scores are computed from this task: spatial and temporal, both using equation (1). For the spatial measure, the error is calculated as the absolute distance between the target pressure position and the measured position of the centre of pressure. The maximum error corresponds to the total distance between the upper and lower pressure targets. The temporal measure’s error is based on how precise in time the change between target positions occurs. This is specifically measured at the time of zero-crossing, respective to the beats of the metronome. Maximum error is the time corresponding to one full period. Both the spatial and temporal absolute errors had skewed distributions; therefore, the median of the error was utilised instead of the mean.

##### Foot tracking task

The subjects controlled the 2D position of a dot on a screen through rotations of their ankle, captured with an inertial measurement unit (IMU) attached to their shoe (see Figure 2B) -- the same setup used as the control interface of the SR3T. The subjects were directed to use ankle rotations only, the result of which they could see as a blue dot on a screen. They had to make the blue dot follow the position of a red dot moving along a figure-of-eight path, as shown in Figure 2B, capturing both the pitch and yaw degrees of freedom of our control interface. The path, the red and blue dots were shown to subjects on a monitor screen in front of them. Each trial is composed of 6 laps around the figure-of-eight path, lasting 35 seconds total. The subjects sat at a height to have their foot freely moving in space (see Figure 2B). The motor-coordination score for this task follows equation (1), with the error defined as the Euclidean distance between the blue and red dots. The maximum error is taken as the maximum recorded error across all time points in all trials of all subjects. Once again, due to the skewness in the absolute error distribution, the median of the error was used in the accuracy calculation.

#### HAMCA hand coordination tasks

##### Hand dimensionality

Subjects performed two toy assembly tasks while wearing a Cyberglove II (CyberGlove Systems LLC, San Jose, CA) to capture the motion of their hand and fingers, with 22 degrees of freedom. The tasks involved assembling a LEGO DUPLO toy train (LEGO 10874), and assembling a toy car (Take Apart, F1 Racing Car Kit) using a toy drill and screws (see Figure 2, D and E). To ensure the appropriate fit of Cyberglove II we made sure all participants had a minimum hand length of 18 cm, measured from the wrist to the tip of the middle finger.

Principal Component Analysis (PCA) was performed on the collected data. We relate a greater number of principal components needed to explain the variance of the motion, to greater hand coordination ^84^. The resulting motor-coordination score is defined as the number of principal components required to explain 99% of the recorded motion’s variance, normalised by the number of degrees of freedom recorded: 22.

##### Piano timing

The subjects used their right-hand index finger to press the same piano key at varying tempi (40bpm, 60bpm, 80bpm, 100bpm and 120bpm) played by a metronome. In total, subjects performed 25 trials in random order (5 at each tempo) composed of 10 keystrokes.

The relevant motor-coordination score follows the same concept as that of equation (1); for further clarity we present it in more detail, in equation (2). The normalised error is the absolute time deviation from the correct tempo divided by its period; that is, the time between keystrokes (inter-onset intervals or IOI) minus the period of each tempo in seconds, as shown in equation (2).

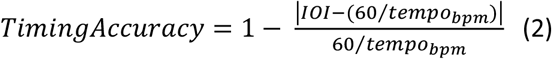

Where tempo is the beats per minute value, hence making 60/tempo the beat period in seconds. Nine samples were generated in each trial (given that nine IOI are generated by ten keystrokes); hence, there were 45 samples generated at each tempo, which had a skewed distribution. The median of these values was taken as the score at each tempo and then the five tempi’s scores were averaged to obtain a single value for their motor-coordination score in the task.

##### Piano jumping

The right-hand index finger was used to move back and forth between two keys and press them at a rate given by the metronome (fixed tempo of 60bpm) – reminiscent of the note and chord jumping technique that pianists train for. Three piano keys were selected, one positioned in the middle of the piano and the other two spaced 7 whole notes to the left and right of it. Three combinations of two keys were to be followed: left and middle, middle and right, left and right (see Figure 2F). In total, subjects performed 15 trials in random order (5 at each key combination) composed of 12 keystrokes each. The relevant motor-coordination score is defined the same way as in the piano timing task. Timings are measured between two consecutive and correct key presses - timings relating to incorrect keypresses were discarded. The latter is done automatically as the note values will be different to what is expected. In order to make up for cases where participants might have pressed the incorrect key, or missed a beat, we consider a window of size of the tempo period (1 second) centred on the correct beat time. If notes are played outside of this window, we assume that the first keystroke of the interval is a wrong one. As the incorrect notes are already removed, a time before the window would mean that the same key was pressed twice consecutively and a time after it would mean that a keystroke was missed. Most of the subjects had no misses or 1 miss out of 165 samples.

##### Piano loudness

On the digital piano, the loudness of a note depends on the velocity with which the relevant key is pressed. A fast press will produce a louder sound and vice versa. This is a skill pianists train for to be able to control the loudness of individual notes and is included as a task-related metric. Subjects were instructed to press a single key at a target level of loudness, with both the target level and the level at which they pressed shown to them visually. Before starting the experiment, participants were instructed on how to set their minimum (0%) and maximum (100%) keystroke loudness values. The piano’s recorded loudness values range from 0 to 127. A very slow key press corresponds to values around 2-8, whereas fast presses fall within values of 120-127. Participants were allowed to familiarise themselves with the visual interface by the experiment runner doing one block of trials on themselves, with the participant watching the interface. Then, they are given up to 5 unrecorded trials to familiarise themselves with how the key presses relate to numerical values, and for the experiment runner to ensure that they cover the full range of values in their key presses. They are then asked each to define their own range, by pressing the key at 0% and 100%. These values are recorded and used to define their range for the experiment. The loudness values for levels 25%, 50% and 75% are obtained by linear interpolation between the 0% and 100% values defined for each participant.

In total, subjects performed 25 trials, 5 at each loudness level (randomised): 0%, 25%, 50%, 75% and 100%, composed of 10 keystrokes each. The motor-coordination score is calculated following equation (1), with the error defined as the deviation from the target values (in percentage loudness) and maximum error as the maximum committed among all the trials of all of the participants for each loudness level (these are as follows, Level 0: 34.0833, Level 25: 42.6250, Level 50: 34.6667, Level 75: 28.4583 and Level 100: 24.4167). After analysing the results, the average motor-coordination score was calculated using only the results at the 25%, 50% and 75% loudness levels given that these targets required more skilled velocity control than the 0% and 100% levels. Thus, their use would enhance individual differences between participants.

#### Piano playing

To assess the participants’ performance on actual piano playing, a sequence with 38 notes played at a constant tempo (isochronous) of 80bpm was devised. Subjects were able to learn and follow the sequence while playing, aided by the software Synthesia (Synthesia LLC). Synthesia showed the notes of the sequence as coloured blocks scrolling on-screen. The participants had to press the keys corresponding to the positions on the keyboard with which the Synthesia blocks were aligned in time to the music in order to score points (see Figure 1). The sequence of notes was designed to be played mainly with the right hand, plus one finger for notes that were too far to the right side of the right hand. These notes could then be reached either using the index finger of the left hand, or, if wearing the SR3T, by activating the robotic finger. On Synthesia, the notes to be played by the right-hand fingers were coloured in green, and the notes to be reached with the extra finger were coloured blue. Similarly, the relevant keys on the keyboard were marked with the same colours (see Figure 2F).

Subjects played the sequence first without and then with the SR3T for 15 trials in each block. The first five trials were considered as practice trials (not recorded), the next ten were recorded but only the last 5 are used for computing the mean piano playing scores per subject due to the fact that subjects were still learning the sequence, especially the ones with no piano playing experience. For the first block of trials, without the SR3T, subjects played using their right hand for green coloured notes while blue coloured ones were played with the left-hand index finger. To achieve this, subjects had to cross their left hand over the right one. For the second block of trials, the left index finger was replaced by the SR3T. We score each individual keypress’s timing as follows:

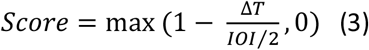

where ΔT is the absolute time difference between the keypress and metronome beat, and IOI is the corresponding beat time period. Therefore, the participants receive a full score for each correct keypress at the exact correct time, with the score linearly decreasing for time deviations, up to half the beat period on each side, at which point the score is 0. Incorrect notes within this window are obviously marked as 0 too. We then average the note scores for the entire sequence.

In some trials, recordings were stopped prematurely due to technical errors. In all such cases, the score is calculated with respect to the recorded section only. However, if less than 50% of the notes are recorded, then the trial is discarded. This happened only in two trials from the same subject which were removed. No other trials for any subjects had any missed recordings. There were also cases where participants missed one initial beat, leading to them being off-beat for the entire sequence. To adjust for this, we calculate the scores for the sequence as originally timed, plus if it were started one beat early, or one beat late. We then take the highest score of the three cases to represent the piano playing score for that trial. This only occurred twice.

#### Statistical Analysis

We first tested for statistically significant differences between the pianists and the naïve players in each of the HAMCA motor-coordination scores, using permutation test. We then calculated the correlation matrix between the motor-coordination scores of all subjects, looking for dependencies between the HAMCA tasks and scores.

For the analysis of the piano playing performance with the LHIF and with the SR3T we addressed trials 5 to 10, after the initial fast learning plateau. Using t test, we looked for significant differences between the pianists and the naïve players in the scores of the trials played with their LHIF and with the SR3T. We then averaged over trials 5 to 10 to get a single piano playing score for each subject in each task to be used in all following analysis. We also merged the two groups (pianists and the naïve players) so that all test of interactions between the HAMCA motor-coordination scores and the piano playing scores were across all subjects. We then calculated the correlations between the subjects’ scores in the piano playing tasks and the HAMCA motor-coordination scores. We further investigated the correlations with Spearman rank correlation scores.

We then fitted linear regression models to explain the piano playing scores using all different sets of HAMCA motor-coordination measures. We fitted models best on single feature, all possible combinations of multiple features, from 2 to all 8. To compare between these models of different complexity we used the corrected Akaike information criterion (AICc) which is modified for small sample sizes.

## Acknowledgments

The authors thank James Cunningham and Anita Hapsari for their contribution to the SR3T platform (see Cunningham et al., 2018).

## Author contributions

AS, PG and AAF did the initial conception of this work; AS, PG, RM and AAF developed the experimental protocol and hardware/software requirements for it; RM ran the experiments; AS, SH, RM, AAF analysed and interpreted the data; AS, SH wrote the manuscript; AS, SH and AAF revised the manuscript.

## Additional information

AS, SH, RM and PG declare no competing financial interests. AAF has consulted for Airbus.

## Data and materials availability

All data needed to evaluate the conclusions in the paper are present in the paper and/or the Supplementary Materials. Our full dataset will be made available on FigShare upon publication. We can also make these available to reviewers upon request.

