## Supplementary figures for "Playing the piano with a robotic third thumb: Assessing constraints of human augmentation"

### Supplementary Materials

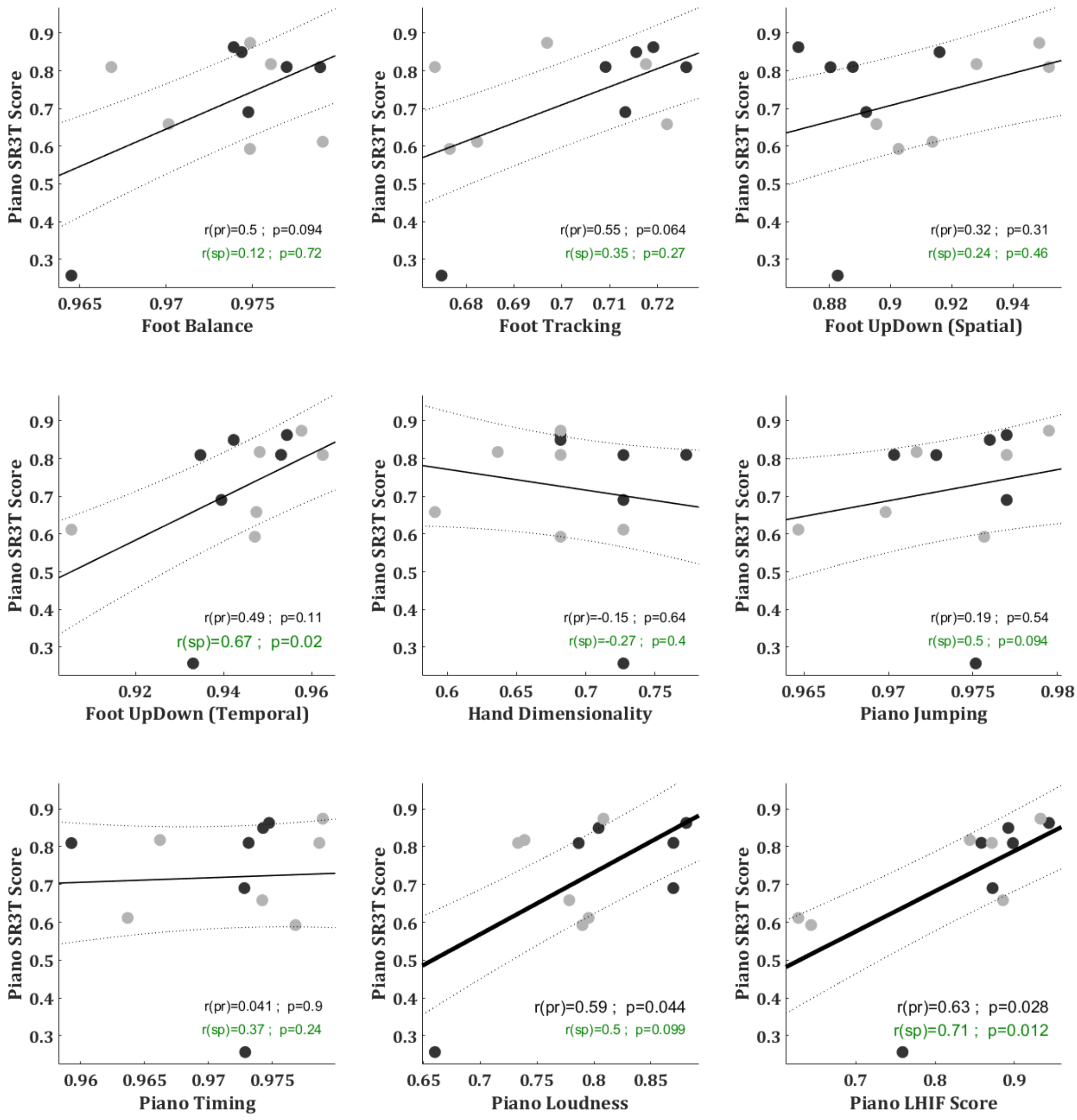


**Supp Figure 1.** ***Correlations between accuracies.*** The first eight panels shows correlations between accuracies in piano playing with the SR3T and in the motor coordination tasks. The ninth panel shows correlations between accuracies in piano playing with and without the SR3T. Naïve subjects marked as grey and experienced subjects as black dots.

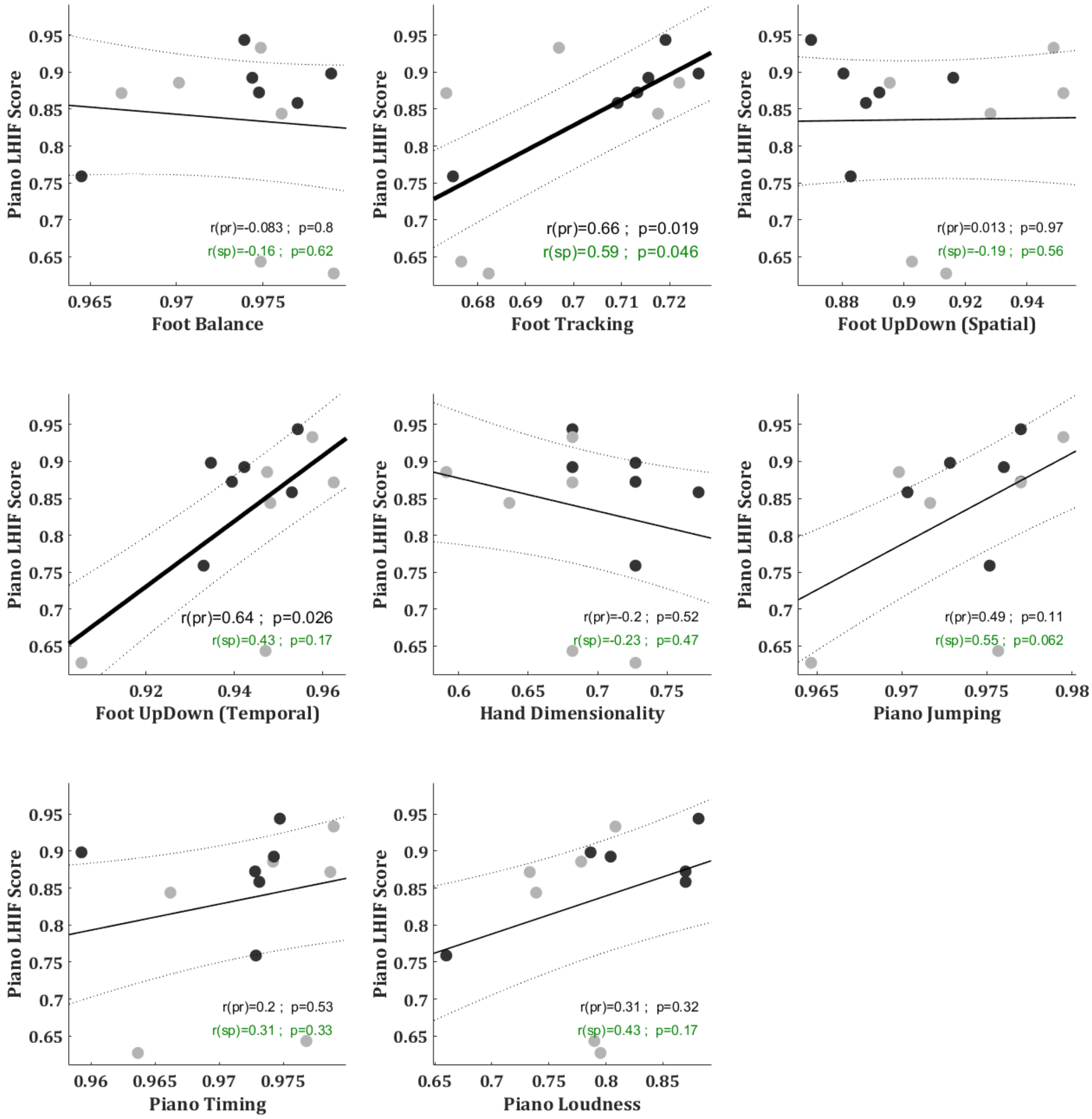


**Supp Figure 2.** ***Correlations between accuracies.*** Correlations between accuracies in piano playing with the LHIF and in the motor coordination tasks.

**Movie S1.** Playing the piano with a robotic third thumb: Assessing constraints of human augmentation, Supplementary Video. Video shows the different experiments described in the manuscript, as well as the supernumerary robotic 3rd thumb (SR3T) in action. It starts with a sequence from our unconstrained pilot experiment, showing our participant after about an hour of practice, using their 10 fingers + the SR3T to play the piano effectively with 11 fingers. This is followed by example videos of our HAMCA setup for motor coordination assessment, as well as our piano sequence task played with the SR3T and the Left-Hand Index Finger (LHIF).
